# Mapping the principal gradient onto the corpus callosum

**DOI:** 10.1101/2020.03.05.978171

**Authors:** Patrick Friedrich, Stephanie J. Forkel, Michel Thiebaut de Schotten

## Abstract

Gradients capture some of the variance of the resting-state functional magnetic resonance imaging (rsfMRI) signal. Amongst these, the principal gradient depicts a functional processing hierarchy that spans from sensory-motor cortices to regions of the default-mode network. While the cortex has been well characterised in terms of gradients little is known about its underlying white matter. For instance, comprehensive mapping of the principal gradient on the largest white matter tract, the corpus callosum, is still missing. We hence mapped the principal gradient onto the callosal midsection using the 7T human connectome project dataset. We further explored how quantitative measures and variability in callosal midsection connectivity relate to the principal gradient values. In so doing, we demonstrated that the extreme values of the principal gradient are located within the callosal genu and the posterior body, have lower connectivity variability but a larger spatial extent along the midsection of the corpus callosum than mid-range values. Our results shed light on the relationship between the brain’s functional hierarchy and the corpus callosum. We further speculate about how these results may bridge the gap between functional hierarchy, brain asymmetries, and evolution.

## Introduction

Cognitive neurosciences assume that cognition is embedded within and constraint by the organization of the brain. The advent of neuroimaging methods granted novel insights into the functional organization of the living human brain. Resting-state functional connectivity (rsfMRI), in particular, has proven useful for mapping multiple functional networks without relying on participants to perform functional tasks in the scanner (Smith et al., 2009; Power et al., 2011; Biswal et al., 1995). rsfMRI networks reveal different time courses and spatially correspond to tasks-related functional activations (Tavor et al., 2016). This spatial correspondence suggests an underlying organization that adheres to fundamental principles. For instance, Mesulam (1998) proposes that brain regions can be organised along a gradient ranging from sensory-motor to higher-order brain processes. In line with Mesulam, most of the variance of rsfMRI resembles a gradient that spans from sensory-motor cortices to regions of the default-mode network (i.e. latter is referred to as the principal gradient, Margulies et al., 2016). While sensory-motor cortices are primary cortices (i.e. idiotypic areas Mesulam, 1998) and the default-mode network has been associated with transmodal higher-order functions (Margulies et al., 2016, Alves et al. 2019). The principal gradient may thus represent one fundamental principle driving a hierarchical organisation of cognitive functions (Huntenburg, Bazin, & Margulies, 2018).

While this principal gradient is putatively the same for both hemispheres (Margulies et al., 2016) little is known on the connections between the left and the right principal gradient. With 200 to 300 millions axons (Tomasch 1954), the largest white matter commissure in the primate brain is the corpus callosum (Burdach 1826). While other routes of interhemispheric interactions are possible (e.g. anterior commissure, Risse et al., 1978), surgically severing callosal connections severaly interrupts interhemispheric functional connectivity in monkeys (Johnston et al., 2008; O’Reilly et al. 2013) and humans (Roland et al., 2017). Concordantly, studies in split-brain patients present with severe behavioral manifestations such as Alien hand syndrome (Akelaitis 1945; Bogen, 1993) Hemispatial Neglect (Kashiwagi *et al.*, 1990; Heilman and Adams, 2003; Lausberg *et al.*, 2003; Park *et al.*, 2006; Tomaiuolo *et al.*, 2010) Optic aphasia (Freund, 1889; Luzzatti *et al.*, 1998) or pure alexia (Dejerine, 1892; Geschwind and Fusillo, 1966) amongst others. The corpus callosum is hence considered critical for the collaborative integration of functions relying on both hemispheres. Despite its crucial role in functions of various hierarchical order, current parcellation approaches divide the corpus callosum based on its cytoarchitecture, geometry and topological projections, with little regard to the callosum functional organization

Several methods have been proposed to parcellate or segment the corpus callosum. Amongst these methods, four main classifications are still currently applied. Post mortem studies identified anatomical features along the midsagittal section of the corpus callosum, including the rostrum, genu, anterior and posterior body, isthmus and splenium (Crosby et al., 1962 ; figure 1A). While this classification is still widely used today, it does not define clear boundaries between the different segments. A second tentatively sharper parcellation was based on histological measures of the midsagittal axonal diameter but it fails to systematically generalise across individuals (Aboitiz & Montiel, 2003, 1992, figure 1B). The enforcement of a geometrical grid of seven parcels has been used to somewhat circumvent the limitations of generalisation (Wittelson, 1989; figure 1C). All these anatomical, histological and geometrical parcellations of the midsection of the corpus callosum do however not consider the functional organisation of the brain areas it connects.

**Figure 1.**
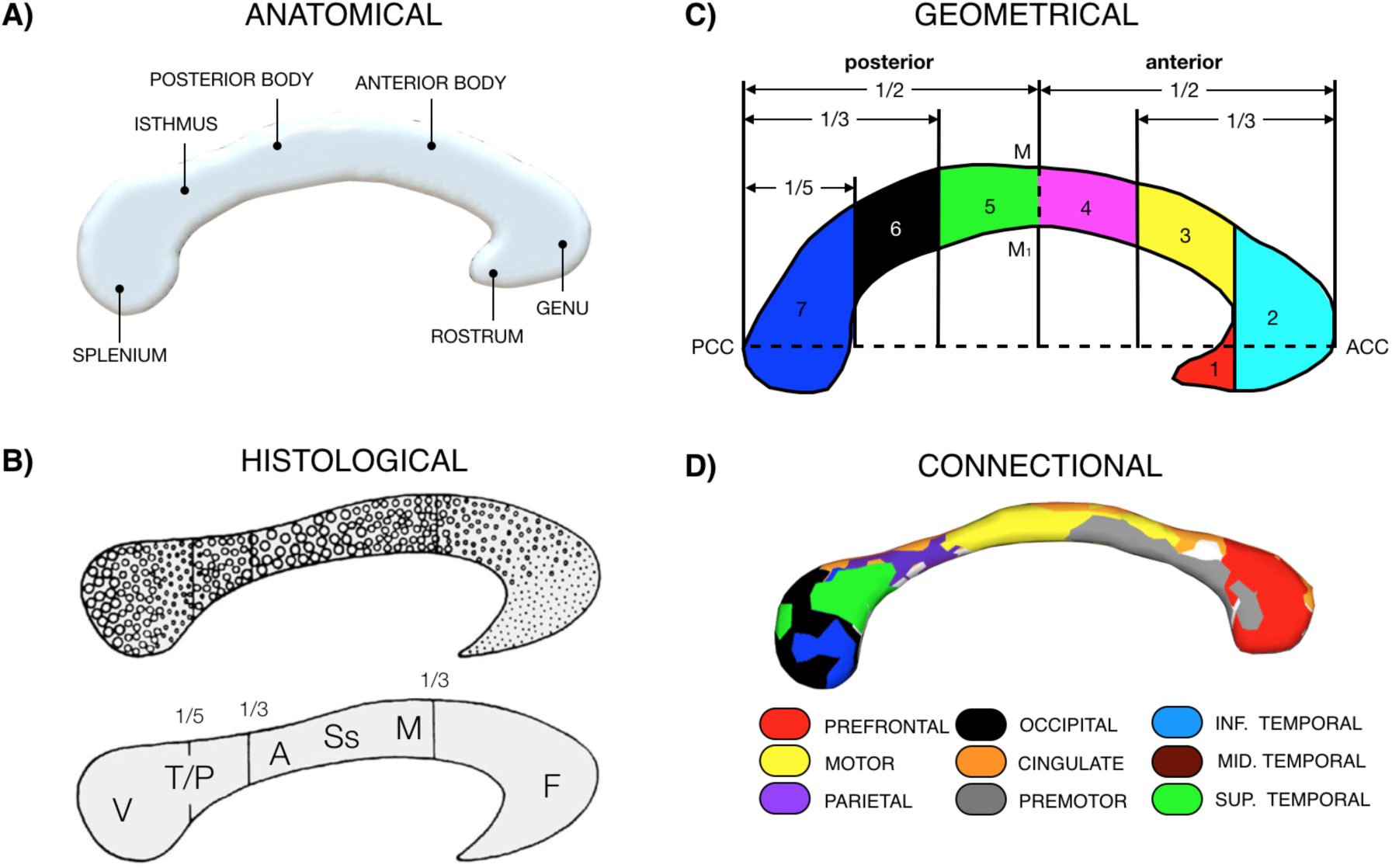
Parcellations of the corpus callosum along the midsection (right view). A) Anatomical division: depicts the anatomical classification (rostrum, genu, anterior and posterior body, isthmus, splenium) based on gross anatomical features. B) Histological division: the dots in this schematic diagram indicate the range of axonal density and associate them with cortical projections to cortical areas (F: Frontal, M: Motor, SS: Somato-Sensory, A: Auditory, T/P: Temporo-Parietal and V: Visual (from Aboitiz & Montiel, 2003). C) Geometrical division: This parcellation generates seven parcels by equally diving the midsection (from Wittelson, 1989). ACC: most anterior point of the corpus callosum, PCC: most posterior point of the corpus callosum, M: middle of the corpus callosum. D: Connectional division: Shows the parcellation of the cortex based on diffusion tractography (from Catani & Thiebaut de Schotten 2012).

Since the functions of the corpus callosum are defined by the areas that it connects to, tract-tracing in non-human primates (Schmahmann and Pandya 2009) or diffusion weighted imaging tractography in humans (Huang et al., 2005; Hofer and Frahm, 2006, figure 1D), is likely the most promising appraoch to investigate the functional organisation of the corpus callosum. However, a comprehensive mapping of the hierarchical organization of cognitive functions on the corpus callosum is still missing. Such a mapping, however, may help to better understand the functional impact of brain lesions that disconnect the corpus callosum.

In this study we, therefore, propose a novel division of the corpus callosum based on the projection of the principal gradient onto the callosal midsection. We also explored how variability in connectivity and quantitative measures of this connectivity relate to the principal gradient values along the callosal midsection.

## Methods

### Dataset

All 7T rsfMRI and diffusion-weighted data used in this study were based on the Human Connectome Project (HCP; Van Essen et al, 2013), which is publicly available and anonymized. For rsfMRI data, we used an averaged map of the principal gradient published in Margulies et al. (2016). This averaged gradient map can be downloaded here: https://neurovault.org/collections/1598/. The minimal preprocessed Diffusion-weighted data consisted of 164 participants (age: mean = 29.44 (3.29); females: 100) from the HCP. Participant recruitment procedures and informed consent forms, including consent to share de-identified data, were previously approved by the Washington University Institutional Review Board

### Gradient percentage maps

We used an averaged map of the principal gradient published (Margulies et al., 2016). In Margulies et al. (2016), functional connectivity matrices were calculated from the HCP dataset and the gradient extracted using nonlinear dimensionality reduction via diffusion embedding (Coifman et al., 2005). In order to assess callosal connections to cortical regions along the principal gradient, we segregated voxels within the cortex based on their location along the principal gradient. We parcellated the principal gradient map into 100 units where each unit represents one percentage of the principal gradient. All operations were conducted using FSL (www.fmrib.ox.ac.uk/fsl). The gradient intensities range between -7.52 and +10.07. A central value of 0 denoted areas not defined along the gradient. For computational reasons, we transposed the gradient values by multiplying the minimum intensity value by -1 and adding the result to all values, thus shifting all values into positive values. We subsequently masked the results with a binarized map of the principal gradient. The resulting image was masked with a cortical grey matter template (i.e. Fast applied to MNI152 template). This resulted in 100 distinct gradient percentage maps, each one presenting a 1% range of the gradient (see figure 2B).

**Figure 2.**
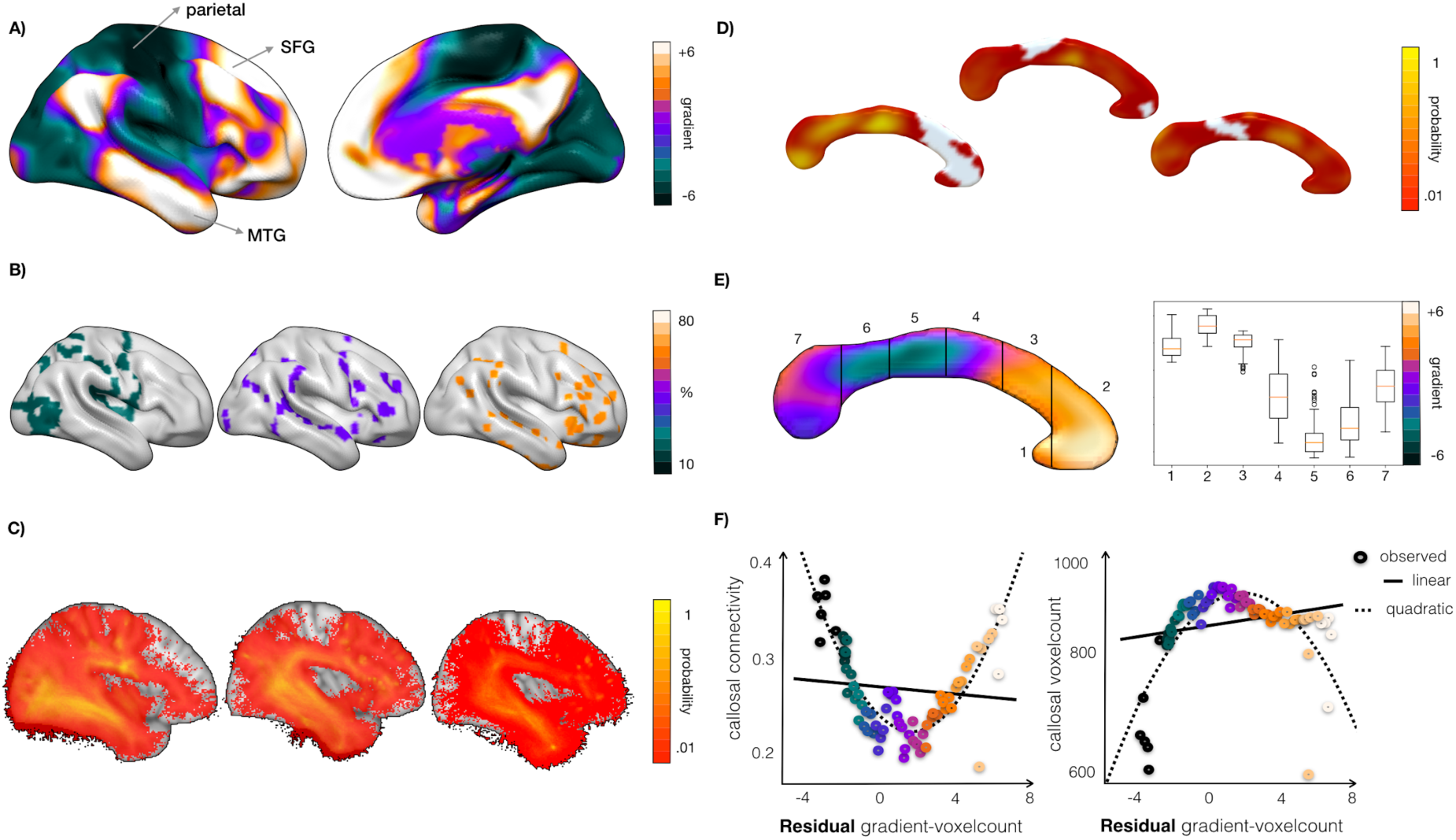
Mapping the principal gradient on the corpus callosum. A) Visualisation of the principal gradient on the cortical surface. B) Examples for percentage maps (20-40-60%) depicting associated cortical areas. C) Percentage-specific tractograms as derived from the percentage gradients maps shown in B. D) Midsagittal display of callosal connectivity based on the tractograms for the 20th, 40th and 60th gradient percentages. The lower the threshold the less participants showed callosal connections in this voxel. E) Callosal gradient map (left), which was computed as the voxel-wise weighted average of callosal connectivity across all percentage maps of the gradient with the Wittelson 7 regions parcellation superimposed. The distribution of gradient values along the Wittelson parcellation scheme is shown on the right. F) Scatterplots showing the relation between callosal connectivity (left) or number of non-zero connectivity voxels (right) and the residual between the gradient corrected for the number of voxels in the respective gradient percentage map. By callosal connectivity, we mean interindividual probability of connection.

### Mapping the principal gradient onto the corpus callosum

In order to map the gradient along the white mater connections, we employed the *Disconnectome* within the BCBtoolkit (Foulon et al. 2018) using a study-specific HCP dataset of 163 participants to generate the underlying white matter tractograms (Karolis et al. 2019). The scanning parameters have been described previously (Vu et al. 2015). In brief, diffusion-weighted imaging consisted of a total of 132 near-axial slices acquired with an acceleration factor of 3 (Moeller et al., 2010), isotropic (1.05 mm3) resolution and whole head coverage with a TE of 71.2 ms and a TR of 7000 ms. These images were acquired with 65 uniformly distributed gradients in multiple Q-space shells (Caruyer et al., 2013) with 6 b0 images. The acquisition was repeated four times with a b-value of 1000 and 2000 s mm−2 in pairs with left-to-right and right-to-left phase-encoding directions. The default HCP preprocessing pipeline (v3.19.0) was applied to the data (Andersson et al., 2012; Sotiropoulos et al., 2013), which consists of susceptibility-induced distortion correction (Andersson et al., 2003), TOPUP (Smith et al., 2004) and subsequent motion and geometrical distortion correction via the EDDY tool implemented in FSL.

For each gradient percentage map, the Disconnectome filtered tractography of the white matter tractograms. The resulting filtered tractography (i.e. streamline passing by the gradient percentage maps) were visually inspected and transformed into visitation maps (i.e. binary mask of the voxels including more that 1 streamline, Thiebaut de Schotten et al., 2011). To generate a group-level visitation map, all normalised individual visitation maps were averaged voxel-by-voxel. This group-level visitation map represents the connection probability per voxel, and thus accounts for the interindividual variability of the anatomical representation of tracts for the respective gradient percentage map (see Figure 2C). All visitation maps are in MNI152 space and are available upon request to the corresponding author. The averaged

This study focussed on the relationship between the functional gradient and the interhemispheric commissural connections of the corpus callosum. We manually delineated the corpus callosum midsection on the midsagittal slice of the MNI152 template. For each gradient percentage, the visitation map was masked with the corpus callosum midsection to reconstruct callosal connectivity maps (see figure 2D). We subsequently created a weighted average map based on the following equation, in which the callosal connectivity maps were divided by the sum of all callosal connectivity maps.

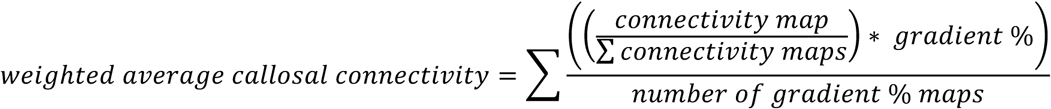

Each of the divided visitation maps was then multiplied by the respective level of the gradient (1% = times 1, 2% = times 2, etc.). This resulted in 100 weighted callosal connectivity maps, one for each gradient percentage. To obtain one single unified callosal connectivity gradient map, the weighted connectome maps were summed up and divided by the number of percentages (n=100). The resulting weighted average connectivity map is shown in Figure 2E. The complete weighted average tractogram is available here: https://neurovault.org/collections/7797/.

### Statistical analyses

Statistical analyses were performed in SPSS (IBM SPSS Statistics, Version 24.0. Armonk, NY: IBM Corp.). In order to avoid spurious measurements, we truncated the distribution of the gradient percentage maps at either end to avoid including maps with less than 300 voxels (supplementary figure 1). Included percentage maps ranged from 13% to 82% (equivalent original gradient values: – 5.23 to +6.90). We explored how variability in connectivity and quantitative measures of callosal connectivity relates to the principal gradient values. The connectivity probability between the corpus callosum and the gradient percentage was calculated by averaging each callosal connectivity map. The callosal space occupied by each percentage was defined as the number of non-zero voxels for each callosal connectivity map. To explore the relationship between the corpus callosum and the principal gradient, we plotted the connection probability against the residuals of the gradient value and the number of voxels of the respective gradient percentage map. To assess the relationship between the value of the gradient and the spatial extent on the corpus callosum unbiased by the number of voxels for each percentage value we regressed out the number of voxels in the scatterplot. The same procedure was applied to plot the callosal space occupied by each percentage. The two distributions were visually inspected and scrutinised with either linear or quadratic regression as applicable.

## Results

### Corpus callosum gradient map

The principal gradient projections on the corpus callosum is displayed below (Figure 2E).

Results indicate a structured distribution of the principal gradient along the midsection of the corpus callosum where the anterior corpus callosum is more likely connected to areas of higher principal gradient values compared to posterior areas. Accordingly, the negative spectrum of the principal gradient peaked in the isthmus and the posterior body, whereas the positive spectrum occupied the anterior part of the body, the genu and the rostrum. In some parts of the splenium a small peak of positive values was also evident.

### Relationship between the principal gradient and callosal connectivity

Visual inspection of the data clearly indicated a U-shaped distribution. Using a curve-fit estimation, we investigated the possibility for either a linear or quadratic relationship between connection probability and the principal gradient. A linear regression model was fitted onto the distribution (Y_i_ = 0.229 + -0.002*X_i_), which did not match the data (F_(1, 68)_ = 0.698, r^2^ = 0.010, p = 0.406). However, both a quadratic regression revealed a highly significant (r^2^ = 0.739, p < 0.001; (Y_i_ = 0.174 + - 0.006*X_i_ + 0.004*X^2^_i_)) relationships between callosal connection probability and the principal gradient as illustrated in Figure 2F.

### Relationship between the principal gradient and associated callosal area

Similar to the previous analysis, a curve-fit estimation was employed to investigate a linear or quadratic relationship between the number of callosal voxels that may connect to a certain gradient value. A linear regression model (Y_i_ = 840.029 + 5.280*X_i_) failed to reach significance (F_(1,68)_ = 3.147, r^2^ = 0.044, p = 0.081), while the quadratic model reached significance (Y_i_ = 906.792 + 10.801*X_i_ + -5.387*X^2^_i_ with F_(2,67)_ = 41.708, r^2^ = 0.555, p < 0.001). The inverted U-shaped distribution of associated callosal areas across the principal gradient is depicted below (Figure 2F).

## Discussion

The current study investigated the relationship between the principal gradient and cortical projections of the corpus callosum. We reveal three main results: First, the apexes of the principal gradient span from the genu to the posterior body. Second, the individual variability of callosal connectivity is lowest at the apexes of the gradient. Lastly, the number of connected voxels in the callosal midsection showed a decrease at both ends of the gradient, while gradient values in the middle appeared more similar. These results will be discussed on the background of evolutionary theories, thus potentially shedding light onto evolutionary processes involved in changes of callosal connectivity.

To depict the principal gradient on the callosal midsection, we mapped each percentage of the gradient independently. The callosal gradient (Figure 2E, left) indicates that the negative spectrum of the principal gradient peaks in the posterior body, whereas the positive spectrum peaks in the genu. Previous research demonstrated that the posterior midbody contains fibers that project to lower-level processing areas (e.g. somatosensory areas; Schmahmann and Pandya 2009). The genu connects to higher-level associative regions (i.e. prefrontal and orbitofrontal cortices; Schmahmann and Pandya 2009). Gradient values in the splenium were heterogeneous with highest values for a region previously shown as connecting the angular gyrus (Park et al., 2008; Chao et al., 2009) well known for its high values in the principal gradient (Margulies et al. 2016) and central structure of the default mode network (Zuo et al, 2012). Hence the spatial layout of the callosal gradient map is concordant with a previously hypothesised anatomical-functional organisation of the corpus callosum (Catani & Thiebaut de Schotten 2012). Gradient values in the different portion of the Wittelson parcellation (Figure 2E, right) indicated a heterogeneous distribution of gradient along the corpus callosum. With some regions (3,2, 1) being highly consistent in terms of gradient values and region 7 to 4 being more distributed. A higher distribution of values might be related to a wider range of neuropsychological syndromes associated with disconnections of the same regions of the corpus callosum (Catani and Thiebaut de Schotten 2012). Additionally, the callosal gradient resembles previous work mapping the diameter and myelination of callosal fibres (Aboitiz et al, 1992; Aboitiz & Montiel 2003). Accordingly, the genu contains poorly myelinated fibers of thin diameter compared to callosal fibers that connect to the posterior midbody. Diameter and myelination are contingent on conduction speed (Hursh, 1939; Waxman & Bennett, 1972). Furthermore, a study comparing conduction delay in various callosal segments showed a fastest speed in unimodal areas (motor, somatosensory and premotor areas) but a longer delay in transmodal temporal, parietal and occipital areas (Caminiti et al., 2013). This would suggest a different interhemispheric conduction speeds of unimodal (i.e. faster) and transmodal areas (i.e. slower). As such, the callosal gradient may putatively be related to different conduction speeds, which needs to be addressed by future research.

There is reason to believe that the general principle demonstrated at the group level might hold on the individual level, given that interindividual variability in callosal connectivity was lowest for the apexes of the gradient. This findings feeds into a recent hypothesis where structural variability is associated with evolutionary stability (Croxson, Forkel, & Thiebaut de Schotten, 2017). The observed callosal connectivity results may thus be linked to the phylogenetic age of cortical areas. Based on the dual origin theory, it has been suggested that multimodal, prefrontal and paralimbic cortices may have evolved in parallel to sensory and motor cortices (Pandy, Petrides, & Cipolloni, 2015). With regards to the gradient architecture, support for this assumption comes from non-human primate studies. For instance, a comparison between resting-state based gradients in human and structural connectivity-based gradients in macaque monkey displayed similar spatial layouts spanning from primary sensory-motor to fronto-parietal regions (Oligschläger et al, 2019). In humans, this transmodal end of the gradient concords with the default mode network that is also present in chimpanzees (Barks, Parr, and Rilling, 2013) and macaque monkeys (Mantini et al., 2011). Hence several pieces of evidence converge towards phylogenetic older apexes of the gradient. While these arguments may support the interpretation that callosal connectivity may be a proxy for evolutionary stability, comparative studies may compare the callosal gradient architecture across primate species and derive phylogenetic models in order to test this assumption.

In addition, more symmetrical functional organization is found for higher probability in callosal connectivity (Karolis et al. 2019) and corresponds to the apexes of the gradient in the present study. This is in line with studies showing that stronger callosal connections are associated with a decrease in perceptual processing asymmetries. The corpus callosum has a long history as an underlying factor of functional hemispheric asymmetries (Ocklenburg et al., 2016) and appears to decrease asymmetries in perceptual processes (e.g. Friedrich et al., 2017). However, this result does not hold true systematically as studies investigating speech processes (i.e.: word fluency and semantic decision making) indicate that ‘stronger’ corpus callosum increase asymmetries between the two hemispheres (Westerhausen et al., 2006, Josse et al., 2008). Hence, the callosal gradient map may be a useful new framework to clarify contrasting results in the role of the corpus callosum in functional hemispheric asymmetries.

A decrease in the variability of callosal connectivity at either end of the gradient was in part accompanied by a decrease of the number of voxels connected in the callosal midsagittal area at both ends of the gradient (Figure 2F, right side). Inversely, callosal connectivity was more variable and involved more voxels in the associated callosal area when distant from the gradient’s apexes. According to the tethering hypothesis, evolutionary expansion of the human cortex may have untethered large portions of the cortex from the constraints of early sensory activation cascades and molecular gradients (Buckner & Krienen, 2013). Similarly, the dual origin suggests that evolution progressively expended two cortical trends, one of which emerges from the hippocampocentric division (Pandya et al. 2015) and forms the limbic system that is cortically represented by the default mode network (Alves et al. 2019). In other words, evolutionary expansion would have created regions less functionally restricted than the apexes of the gradient. This, in turn, could lead to a more diverse pattern of connectivity (i.e. higher number of voxels in the associated callosal area) which is in accordance with the observed higher individual variability and the tethering as well as the dual origin hypotheses.

Mapping the principal gradient onto the corpus callosum grants insight into the distribution of callosal connections across functions of various hierarchical levels. Contrary to categorical segmentation methods, our approach visualizes a transient gradient of areas in the callosal midsection based on the connectivity of unimodal and transmodal cortical areas. Therefore, this map may guide clinical investigation of brain lesions that disconnect the corpus callosum with a novel gradient-based perspective. It is important to stress that the current work is not a new parcellation of the CC. In the current work we aimed to avoid a categorical parcellation in favour of a map that shows smooth transitions of gradient with high and low values. As we work in the group average space, a continuum of values would be more accurate than a categorical parcellation. Future research might explore the variability of division of the corpus callosum using gradients values and advanced tractography at the individual level. However, some limitations need to be noted. While we describe the first corpus callosum principal gradient map, this map depicts the weighted average of structural connectivity, based on an averaged map of the principal gradient. The weighted average of structural connectivity allowed us to assess - to some extent-the variability of callosal connectivity, however, individual patterns of the principal gradient are yet unavailable. Therefore, the effect of individual variability of the principal gradient’s spatial layout on callosal projections could not be assessed in this study. Addressing the extent to which the principal gradient and its callosal connections differ across individuals for example via analysis platforms (e.g. Vos de Wael et al., 2020) will be a matter of future research.

Another limitation comes from the chosen tractography approach. Tractography analyses can produce inaccurate results (Jones and Cercignani, 2010; Maier-Hein et al., 2017). To avoid these caveats, we employed methods that have previously demonstrated high anatomical reliability when compared to axonal tracing (Thiebaut de Schotten et al. 2011a; 2012) and with post-mortem dissections (Catani et al., 2012; Thiebaut de Schotten et al., 2011; Catani et al., 2017; Vergani et al, 2014). In the disconnectome tool, all image matrices were binarized depending on the existence of absence of streamlines connecting the corpus callosum with a given gradient percentage mask. In line with other studies (Gong et al., 2009; Shu et al., 2011), we chose a binarization threshold of 1 because the number of streamlines does not reflect the connectivity strength or the true number of axonal projections between two brain regions (Gong et al., 2009; Jones et al., 2013) and previous work shows that changing streamline count threshold for binarization does not change the overall results of network connectivity analyses (Shu et al., 2011). However, this approach does not prevent that each streamline may cross a voxel of different gradient values, leading to potential overlap, especially if gradient values are partially asymmetric. The current approach does not allow to assess gradient asymmetries because of methodological limitations inherent to the gradient estimation (including for instance asymmetries of the seeding regions, asymmetrical template, contamination of the functional connectivity by the other hemisphere, white matter asymmetries). Future studies may wish to circumvent all these limitations to investigate hemispheric asymmetries of the principal gradient along the callosal connectivity.

Futuremore the current approach studies may wish to quantify this problem potentially to investigate hemispheric asymmetries in the callosal connectivity along the principal gradient.

## Conclusion

Our study provides the first comprehensive map of the callosal cross-section that displays the distribution of callosal connections according to a hierarchical organisation of cognitive functions, without enforced boundaries. Our findings align with previous considerations about the evolutionary changes in the brain. Future studies may utilize the map of the callosal gradient to bridge the gap between brain asymmetries and functional hierarchy.

## Acknowledgements

European Research Council (ERC) under the European Union’s Horizon 2020 research and innovation programme (grant no. 818521)

## Notes

### Competing Interest Statement

The authors have declared no competing interest.

### Summary of Updates

1. We changed the wording of some sections for clarity. 2. The central result figure was modified to show the results of an added description of how the gradient values differ between callosal segments defined by the Witelson Parcellation. 3. we added a paragraph in the discussion that links the callosal gradient map to brain asymmetries. 4. We extended the limitation section 5. Some technical information was added in the method section.

