## supplementary figure 1 for "Mapping the principal gradient onto the corpus callosum"

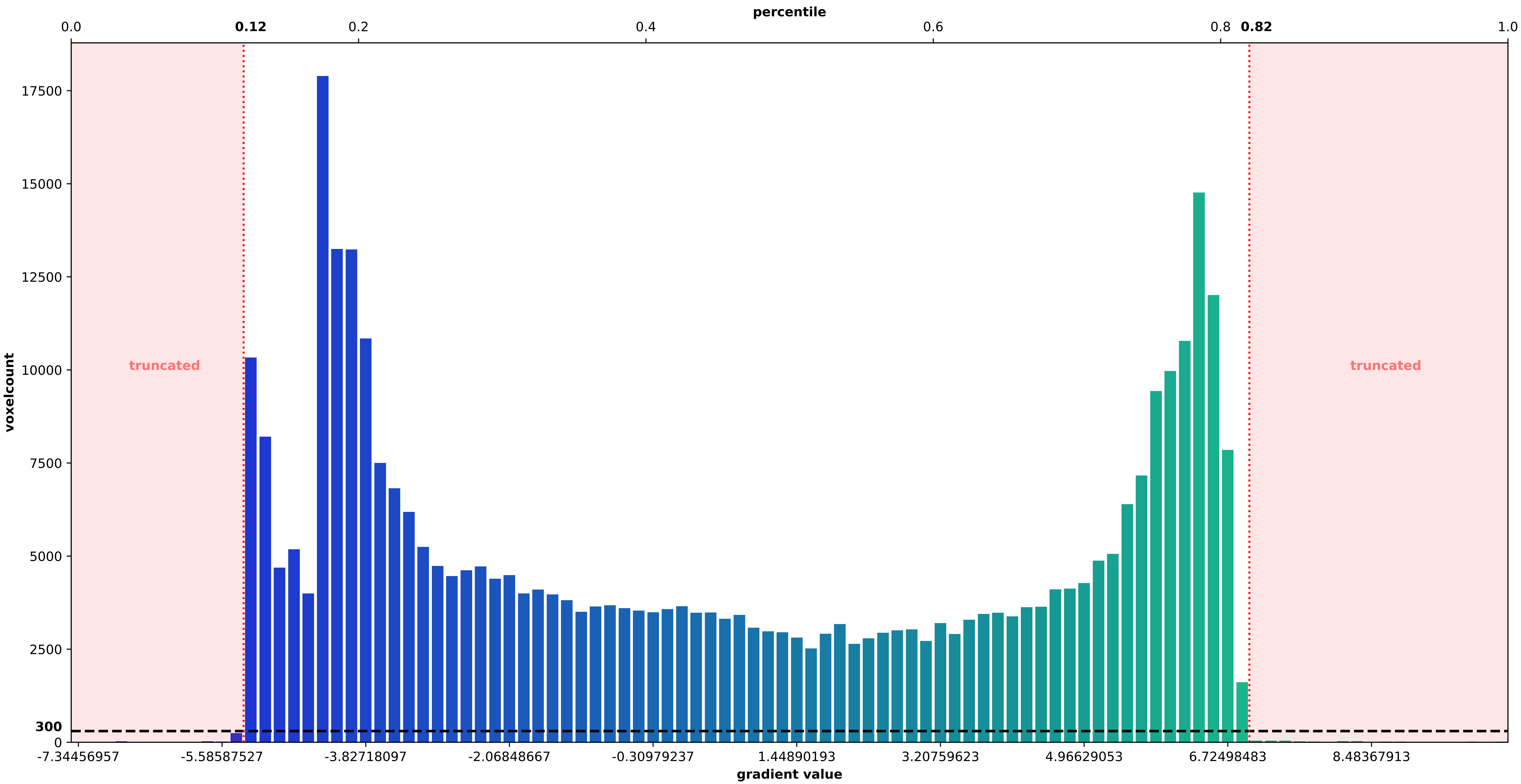


Figure S1. Voxelcount per gradient value. Gradient percentile maps with less then 300 voxels were truncated from the analyses to avaid spurious measurements
